# Organic mulching alters ground-dwelling arthropod communities and promotes key predators in potato agroecosystems

**DOI:** 10.1101/2025.07.14.664685

**Authors:** Julian Winkler, Simeon Leisch, Sascha M. Kirchner

## Abstract

Widespread declines in insect biomass and biodiversity have raised concerns about the sustainability of ecosystem services in agroecosystems. Ground-dwelling generalist predators such as carabids (Carabidae) and spiders (Araneae) are key contributors to natural pest control, yet their populations are threatened by modern farming practices. Organic mulching – a practice that increases within-crop habitat diversity – has been proposed as a means to support beneficial arthropod communities while also delivering agronomic benefits.

This study investigated the effects of different organic mulches (triticale/vetch, straw, grass silage) on carabid and spider communities in three large-scale, replicated field trials in potato fields in central Germany. Arthropods were sampled using pitfall traps during and after the growing season, with additional traps used to assess potential sampling biases related to mulch presence. Community composition, species-specific responses, and potential carry-over effects into the following year were analysed using a combination of generalised linear mixed models, diversity indices, ordination methods, and indicator species analysis.

Across the trials, 12,076 adult carabids (42 taxa) and 2,399 spiders (38 taxa) were collected, enabling robust analysis of treatment effects on community composition and species-specific responses. The results showed that organic mulching, particularly with triticale/vetch, significantly increased the abundance of several carabid (*Bembidion lampros, B. quadrimaculatum, Poecilus cupreus*) and Linyphiidae spider (*Erigone atra, E. dentipalpis, Agyneta rurestris*) species, and altered overall community composition. Species richness tended to be higher in mulched plots, though effects on diversity indices were less consistent. However, positive impacts on predator populations were largely restricted to the potato growing season, with no evidence of persistent carry-over effects into the subsequent year. The study discusses habitat complexity, microclimate moderation, and changes in prey availability as likely mechanisms underlying these patterns.

## 1 Introduction

Over the past three decades, a substantial decline in insect biomass has been well documented (Hallmann et al., 2017; Ziesche et al., 2023), with factors such as habitat change, pesticide use, biological invasions, and climate change identified as primary drivers of this trend (Sánchez-Bayo & Wyckhuys, 2019). Among the affected groups, carabid beetles have also experienced biodiversity losses (Homburg et al., 2019; Skarbek et al., 2021), though these declines are generally less pronounced compared to other insect taxa (Sánchez-Bayo & Wyckhuys, 2019). Nevertheless, recent studies indicate that the widely used pyrethroid insecticides can virtually eliminate many typical carabid and spider species from agricultural landscapes, highlighting the vulnerability of these taxa to modern agricultural practices (Blanco-Moreno et al., 2024).

Ground-dwelling generalist predators, particularly carabids (Carabidae) and spiders (Araneae), play a crucial role in maintaining ecosystem services within agricultural systems. Their ecological importance is underscored by numerous studies demonstrating their effectiveness in suppressing aphid populations, thereby contributing to natural pest control (Diehl et al., 2013). Beyond aphids, carabids are known to prey on slugs (Oberholzer & Frank, 2003; Symondson, Glen, et al., 2002) and weed seeds (Carbonne et al., 2020; Honek et al., 2013), while spiders consume a broad spectrum of arthropods, including several significant crop pests (Kuusk et al., 2008; Saqib et al., 2021). Although generalist predators are regarded as less efficient than specialist predators in controlling specific pests (Diehl et al., 2013), they offer distinct advantages: Generalists typically exhibit greater temporal persistence in agroecosystems and can respond rapidly to sudden increases in pest populations, thereby providing a more stable and resilient form of biological control over time (Symondson, Sunderland, et al., 2002). Thus, preserving their populations is essential not only for biodiversity but also for the continued provision of vital ecosystem services in agricultural landscapes.

Increasing within-crop habitat diversity has been shown to benefit arthropod predator populations (Gurr et al., 2003, 2017). For example, Sunderland & Samu (2000)reviewed evidence that practices such as undersowing and organic mulching often lead to higher spider abundance. Organic mulching refers to the application of plant material, like straw or other crop residues, to the soil surface. In potato cultivation, mulch is typically applied just before the plants emerge; cereal straw has been commonly used in previous studies, though any freshly cut or preserved plant material could be used (Alyokhin et al., 2019). Organic mulching offers several agronomic and ecological benefits. It reduces evaporation (Edwards et al., 2000; Johnson et al., 2004), soil erosion (Döring et al., 2005; Edwards et al., 2000), and can increase crop yields, making mulching an economically viable practice (Wenzel et al., 2024; Winkler et al., 2024).

Organic mulching has been demonstrated to reduce populations of several important agricultural pests. For example, it can lower aphid infestations and the transmission of aphid-borne viruses (Kennedy et al., 2010; Kirchner et al., 2014; Winkler et al., 2025), as well as suppress cabbage stem flea beetles (*Psylliodes chrysocephala*) (Seimandi-Corda et al., 2023), and Colorado potato beetles (*Leptinotarsa decemlineata*) (Genger et al., 2018; Weiler et al., 2025a; Winkler et al., 2024). The reduction in aphids and virus incidence is partly attributed to disrupted host-finding based on visual cues (Döring et al., 2004; Döring & Kirchner, 2022). Colorado potato beetles avoid colonising mulched potatoes and their development appears to be negatively affected by the altered microclimate (Weiler et al., 2025a, 2025b).

However, much of this research was conducted outside Central Europe or identified natural enemies only to the family level, limiting the transferability of the results. Most studies also relied on small plot sizes and did not investigate carry-over effects beyond the main growing season, underscoring the need for more regionally adapted and detailed research.

While the mechanisms underlying the suppression of Colorado potato beetles and aphids are becoming clearer, less is known about how organic mulching affects cabbage stem flea beetles. Some research suggests that reductions in pest populations may be associated with increases in natural enemy populations, particularly carabid beetles and spiders, within mulched fields (Brust, 1994; Johnson et al., 2004; Rämert, 1996; Schmidt et al., 2004). Despite these promising findings, several factors limit the scope of current research. Notably, about half of these studies were conducted on the American continent, where arthropod communities differ markedly from those in Europe. Moreover, most studies exhibited coarse taxonomic resolution, particularly for spiders. Additionally, methodological constraints such as small experimental plot sizes, a lack of comprehensive biodiversity assessments, and the absence of data on carry-over effects beyond the main growing season further restrict the robustness of the conclusions.

To address these gaps, we conducted three large-scale field trials in potato fields in central Germany. Ground-dwelling predators were sampled using pitfall traps during and after the growing season, allowing us to examine both species-specific responses and changes in overall community structure. We also set additional traps in mulched plots with the surrounding mulch cleared away to check whether the immediate presence of mulch biased arthropod activity. Our research objectives were: (i) to assess the effects of different organic mulches on carabid and spider abundances and community composition; and (ii) to investigate potential carry-over effects of mulching on arthropod abundance beyond the main growing season.

## 2 Materials and methods

### 2.1 Field experiments

This study comprised three field experiments: two in 2021 (referred to as trial 2021a and 2021b) and one in 2022. The trials were conducted on commercial organic potato fields near Göttingen, central Germany. A randomized complete block design with three or four replicates (blocks) was used to compare unmulched control plots with plots mulched with triticale/vetch, straw, or grass silage. Each plot measured approximately 500 m² (18 m x 28 m). Organic mulches were applied around the time of potato emergence (June 4, June 11, and May 19/20 for trials 2021a, 2021b, and 2022, respectively). Mulching rates were 35–60 t/ha fresh matter (∼25% dry matter) for triticale/vetch and grass silage, and 4–6 t/ha for straw (∼90% dry matter). The triticale-vetch-mixture was chopped to about 5–7 cm length and the straw was of a similar length. Weather data were obtained from a (Deutscher Wetterdienst, 2025) station (2025) in Göttingen using the *R* package `rdwd` (Boessenkool, 2023). Yields were assessed by harvesting four rows of 3 m in each plot. Further details on the trial setup and management can be found in (Winkler et al., 2024).

### 2.2 Arthropod sampling

Generalist predators (spiders and carabids) were monitored using pitfall traps, following the recommendations of Hohbein & Conway (2018). Each trap consisted of two nested 500-ml transparent plastic cups (opening diameter 105 mm). No covers or guiding fences were used. Traps were filled with 300 ml of tap water containing an odorless detergent and placed on the potato ridges. The holes for the traps were dug just before the first sampling interval, ensuring the rims were level with the surrounding ground.

Sampling occurred in June/July for all trials, again in late August/early September (before harvest) in trials 2021a and 2021b, and around late October (after harvest) in trial 2022. In addition, as a follow-up to trials 2021a and 2021b, carabids were sampled in July 2022 to investigate carry-over effects of mulching in the following year. At that time, trial 2021a was planted with squash (block I and II) and faba bean (block III and IV), and trial 2021b with a grass-clover mixture. Sampling periods are shown in Figure 2. Traps were emptied every 2–5 days during sampling (mostly every 3–4 days). Exact dates are given in Appendix 1. Damaged traps were not emptied but discarded, and their catch counts were recorded as “NA”. When traps were emptied, they were cleaned and refilled with fresh water and detergent. Arthropods were collected using a fine sieve (0.5 mm mesh) and transferred to plastic containers containing 70% ethanol for preservation.

In all plots, two traps (type ‘a’) were placed 6 m from the plot edges (Figure 1). During the June/July sampling in the triticale/ vetch plots of trials 2021a and 2021b and the straw plots of trial 2021b, two additional traps (‘b’) were set in the lower left and upper right centers of each plot. Around each ‘b’ trap, a 1-m diameter area of mulch was removed. Comparing catches between trap types ‘a’ (with mulch) and ‘b’ (mulch removed) was intended to reveal any bias due to the immediate presence of mulch. This approach follows Greenslade (1964) and Melbourne (1999), who compared traps in vegetation to traps with cleared surroundings.

**Figure 1:**
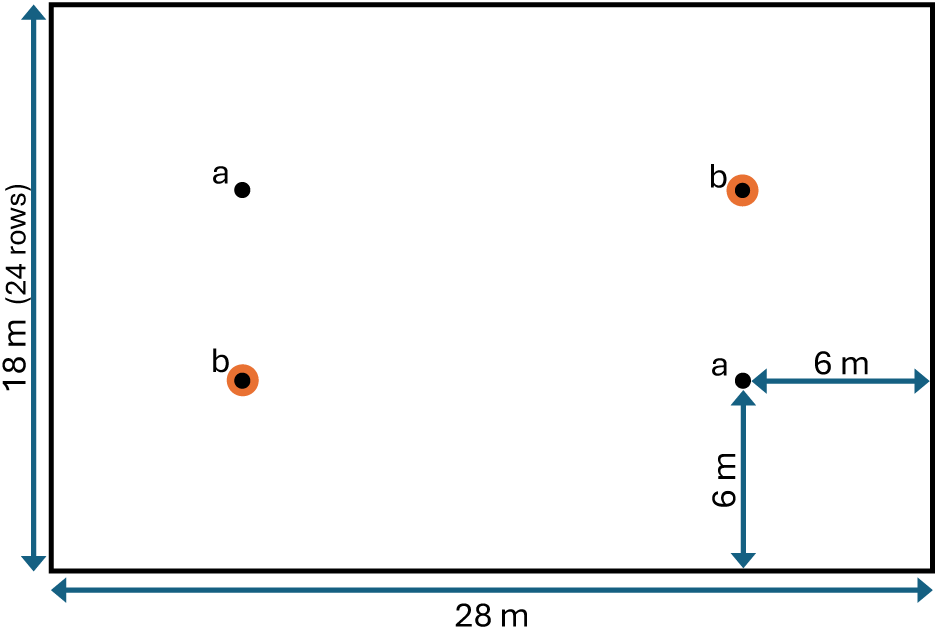
Arrangement of pitfall traps within the plots. a: traps that were placed in all plots. b: traps placed only in triticale/vetch mulched plots (trial 2021a and 2021b) and straw mulched plots (trial 2021b).

**Figure 2:**
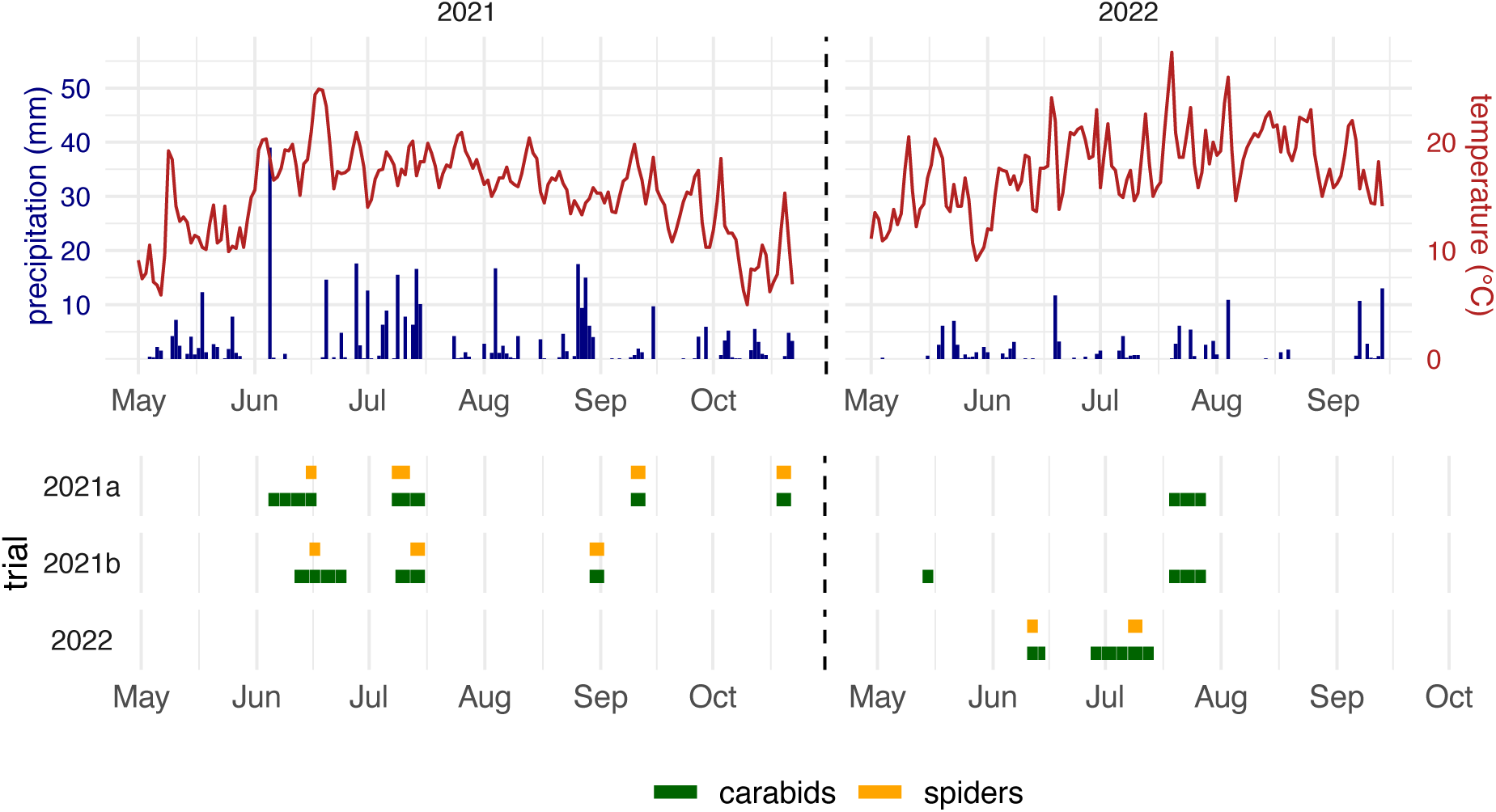
Daily precipitation, mean daily temperature (top graph) and sampling/identification periods of trials 2021a, 2021b and 2022 for carabids and spiders (bottom graph).

### 2.3 Arthropod identification and data processing

All captured arthropods were identified in the laboratory using a stereomicroscope, with magnifications up to 40× for carabids and 80× for spiders. Standard identification keys were used (Freude, 2004; Lindroth, 1974; Lompe, 2025; Nentwig et al., 2025; Roberts, 1995). Approximately 300 spider individuals remained unidentified as they were either destroyed, immature, or too infrequent to allow for reliable identification. Carabid and spider abundance data were analysed using *R*, version 4.4.1 (R Core Team, 2024) and *RStudio*, version 2024.09.0 (Posit Software, PBC, 2024). Raw count data from all pitfall trap types within each plot – including traps with and without mulch removal – were used to calculate descriptive statistics. Data preparation was done using the *R* packages `tidyverse` and `tibble` (Müller & Wickham, 2023; Wickham et al., 2019). Plot means and standard errors were calculated for visualization. For graphical comparison of trap types, means were calculated separately for traps with mulch and those with mulch removed.

For interferential comparisons of carabid and spider abundance between treatments, generalized linear mixed models were fitted to data aggregated by block using the `glmmTMB` package (Brooks et al., 2017). Model diagnostics were conducted with the `DHARMa` package (Hartig, 2024). To compare treatments, Dunnett’s post-hoc tests were performed using the `multcomp` package (Hothorn et al., 2008). All plots were generated with `ggplot2` (Wickham, 2016); supplemented by the `lubridate`, `ggpubr` and `ggh4x` packages (Brand, 2024; Grolemund & Wickham, 2011; Kassambara, 2023).

The community ecology *R* package `vegan` (Oksanen et al., 2024) was used to calculate diversity measures and perform ordination methods. These calculations were performed on each trial’s June/July sampling data. For plots with n = 2 traps per plot, pitfall trap catches in June/July were summed up for individual pitfall traps. For plots with n = 4 traps per plot (triticale/vetch in trials 2021a and 2021b, straw in trial 2021b), means were calculated from one trap with a mulched and one with a cleared surrounding. This gave two data points per plot used to calculate the species richness and evenness. Data were visualised with horizontal jitter plots for each trial, showing individual trap values and mean indicators.

To identify species significantly associated with specific treatments, we performed a multi-level pattern analysis (Dufrêne & Legendre, 1997) using the `multipatt` function in the `indicspecies` *R* package (Cáceres & Legendre, 2009). This analysis was conducted with 999 permutations and a significance level of 0.05. We constructed a Bray-Curtis dissimilarity matrix from the abundance data for non-metric multidimensional scaling (NMDS) using the `metaMDS` function (`vegan` package). Treatment centroids were calculated and plotted on the NMDS. PERMANOVA tests (using `adonis2` in `vegan`) were performed on the dissimilarity matrices to assess treatment effects on community composition, using 999 permutations and stratifying by block. After confirming significant treatment effects and verifying homogeneity of dispersion (with the `betadisper` function), we conducted pairwise PERMANOVAs with Bonferroni correction for multiple comparisons.

## 3 Results

A total of 12,076 adult carabid beetles, representing 42 taxa, and 2,399 spiders from 38 taxa were collected and identified across the three field trials. Mulch treatments significantly affected both the abundance and community composition of these arthropod groups, with more pronounced effects observed for carabids than for spiders. In particular, the triticale/vetch mulch treatment led to a substantial increase in carabid beetle abundance in trials 2021a and 2021b (both p < 0.001, GLMM, α = 0.05), whereas no significant effect was detected in trial 2022. Similarly, spider abundance was significantly higher in triticale/vetch plots in trials 2021a (p < 0.001) and 2022 (p = 0.014), but no effect was observed in trial 2021b.

### 3.1 Weather

Distinct differences in temperature and precipitation characterised the sampling periods of 2021 and 2022. June 2021 featured unusually high temperatures during the sampling period of trial 2021b. Trial 2021a began with a heavy rainfall event (39 mm) on June 5, followed by an extended dry period. Late June to mid-July saw moderate rainfall during July sampling in both 2021 trials. In contrast, 2022 had higher average temperatures than 2021, with exceptionally high temperatures in July during the second sampling period. Notably, 2022 experienced unusually low rainfall throughout all sampling periods.

### 3.2 Carabid response

Carabid catches showed distinct temporal fluctuations across all three field trials during June/July (Figure 3). The triticale/vetch treatment substantially increased carabid abundance in trials 2021a and 2021b (2.3 and 1.6 times higher than control, respectively) but showed slight reduction in trial 2022 (0.9 times control). The straw treatment in 2021 trials resulted in abundances similar to or slightly higher than control (1.0 and 1.2 times), while the silage treatment in 2022 showed slightly reduced abundance (0.9 times control).

**Figure 3:**
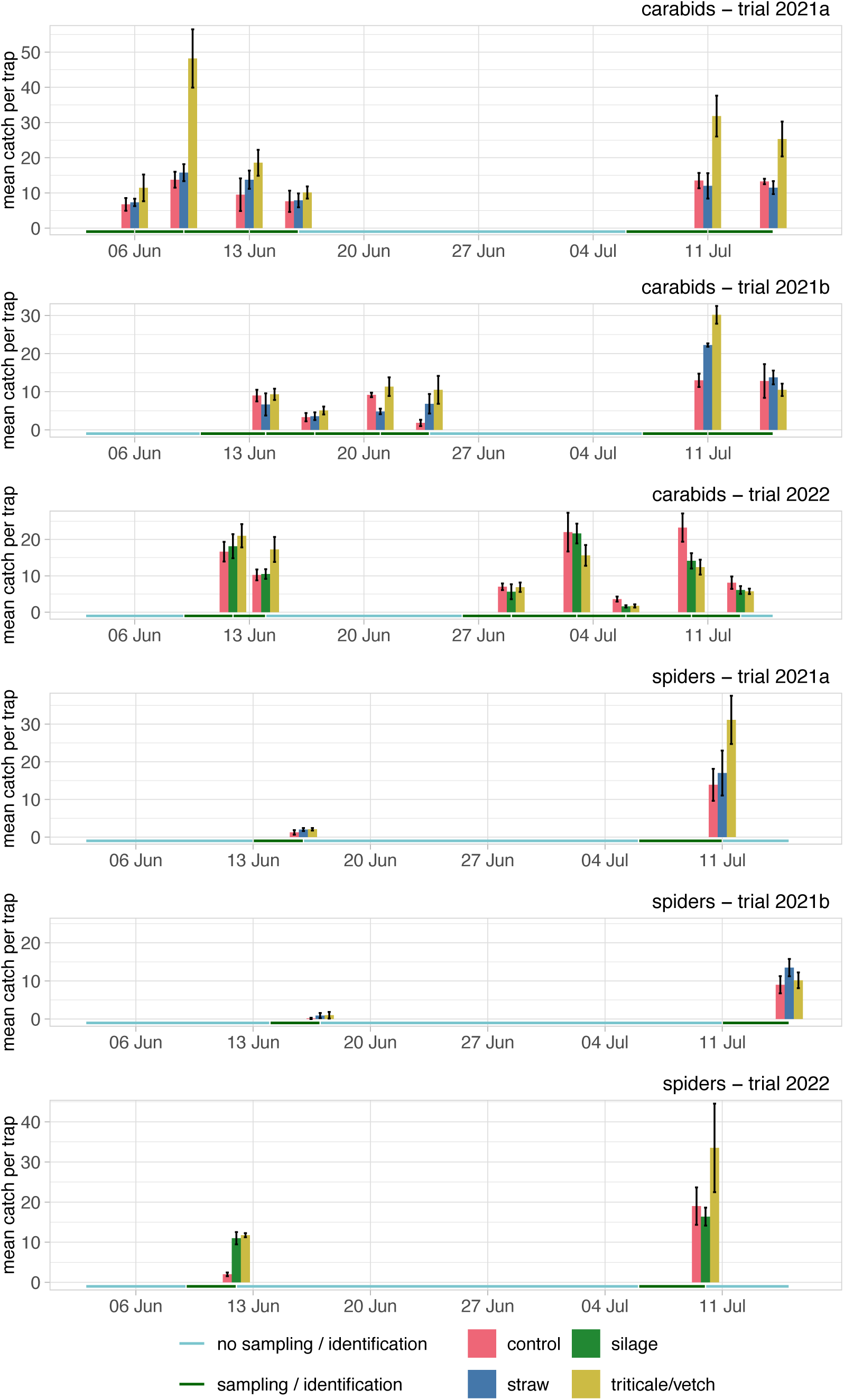
Mean total catches (± SE) of all carabids and spiders in pitfall traps during June/July. Values represent means calculated from plot means (n = 4 in trials 2021a and 2022, n = 3 in trial 2021b). The green horizontal bars indicate sampling periods for the specific catch. See also Appendix 1 for the exact dates and days of sampling.

#### 3.2.1 Species-specific response

The most frequent species found were, in descending order, *Harpalus rufipes*, *Bembidion lampros*, *Poecilus cupreus*, *Bembidion quadrimaculatum*, *Anchomenus dorsalis*, and *Pterostichus melanarius* (see Appendix 2 for a detailed list). Species composition varied considerably among trials (Figure 4). Trial 2021a was dominated by *P. cupreus*, trial 2021b by *Bembidion* species, and trial 2022 by *B. lampros*. Several species showed consistent responses to treatments:

**Figure 4:**
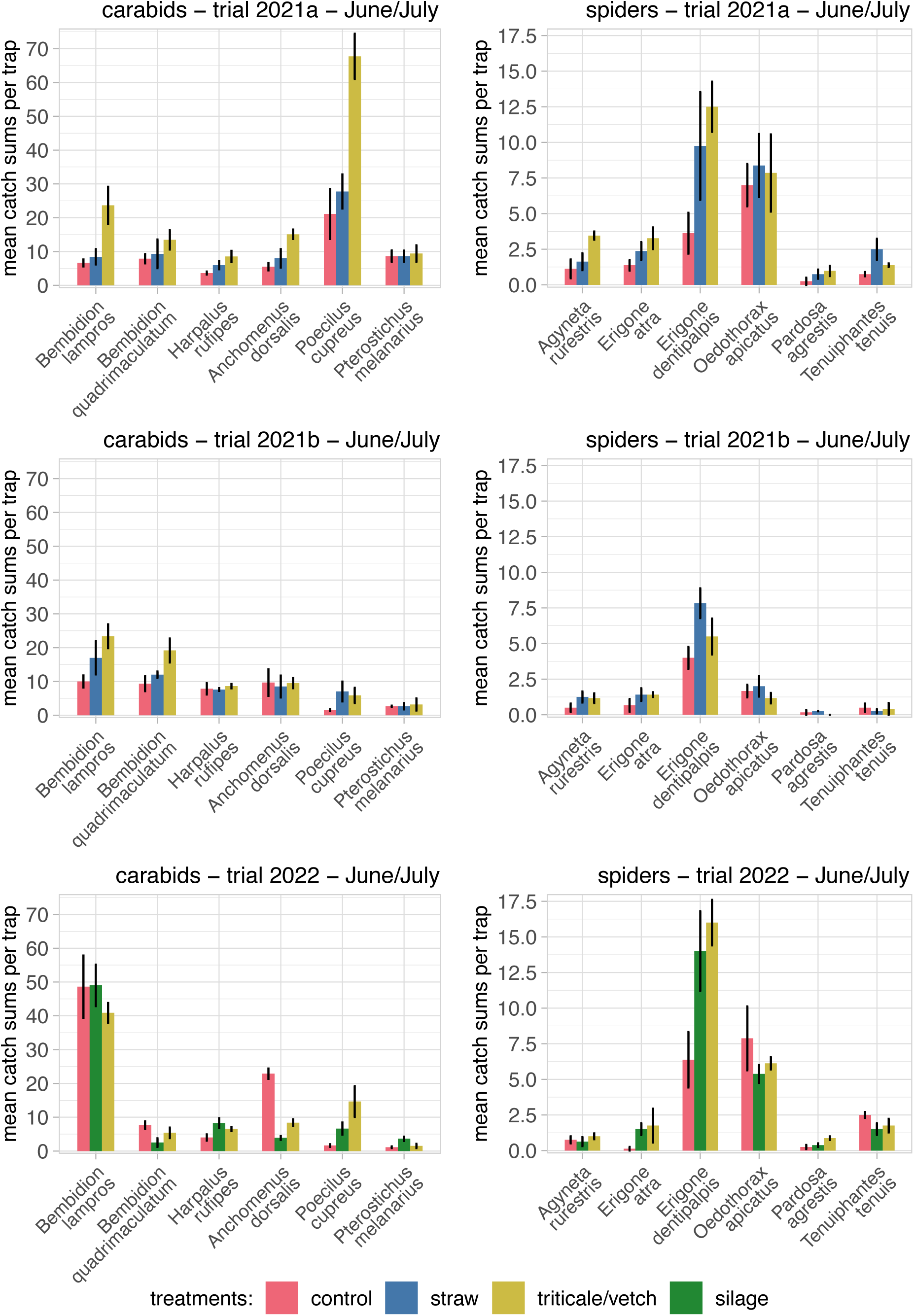
Mean catches (± SE) of the six most frequent carabid species in June/July summed up from a total of 22 sampling days each. Values represent means and SE calculated from plot means (n = 4 in trial 2021a and 2022, n = 3 in trial 2021b).

- *P. cupreus* showed the strongest positive response to triticale/vetch treatment, with 3.3-9.0 times higher abundance compared to control plots across all trials. This species was also more abundant in straw/silage treatments (1.2–4.4 times control) and was mainly active during June.

- *Bembidion* species responded positively to triticale/vetch in trials 2021a and 2021b (2.6 and 2.2 times control) but showed reduced abundance in trial 2022 (0.8 times control). Their response to straw was more modest (1.3–1.5 times higher).

- *A. dorsalis* showed similar frequencies across treatments in trials 2021a and 2021b but was more abundant in control plots in trial 2022. This species was found almost exclusively in July in 2021 trials but more often in June in trial 2022.

The multi-level pattern analysis confirmed significant associations between *P. cupreus* and the triticale/vetch treatment in trials 2021a and 2022 (p=0.023 and p=0.022), and between *B. lampros* and triticale/vetch in trial 2021a (p=0.022). In trial 2022, *A. dorsalis* and *Pt. melanarius* were significantly associated with the control (p=0.022) and silage treatment (p=0.022), respectively.

#### 3.2.2 Carry-over eDects

Late-season sampling (August-October) showed substantially lower carabid abundance, with *H. rufipes* dominating in late August/early September and *Nebria brevicollis* prevailing in October (Appendix 3). In trial 2021a, catches of H. rufipes were higher in the mulch treatments until September, but were slightly lower in trial 2021b.

Analysis of carabid beetle catches from July in the year following mulch application (2022 for trials 2021a and 2021b) revealed no persistent effects of the previous year’s treatments on carabid abundances (Figure 5). *H. rufipes* dominated the carabid assemblages in the follow-up sampling of trials 2021a and 2021b, respectively. Secondary species included *Pt. melanarius* in the follow-up to trial 2021a and *B. lampros* in the follow-up to trial 2021b.

**Figure 5:**
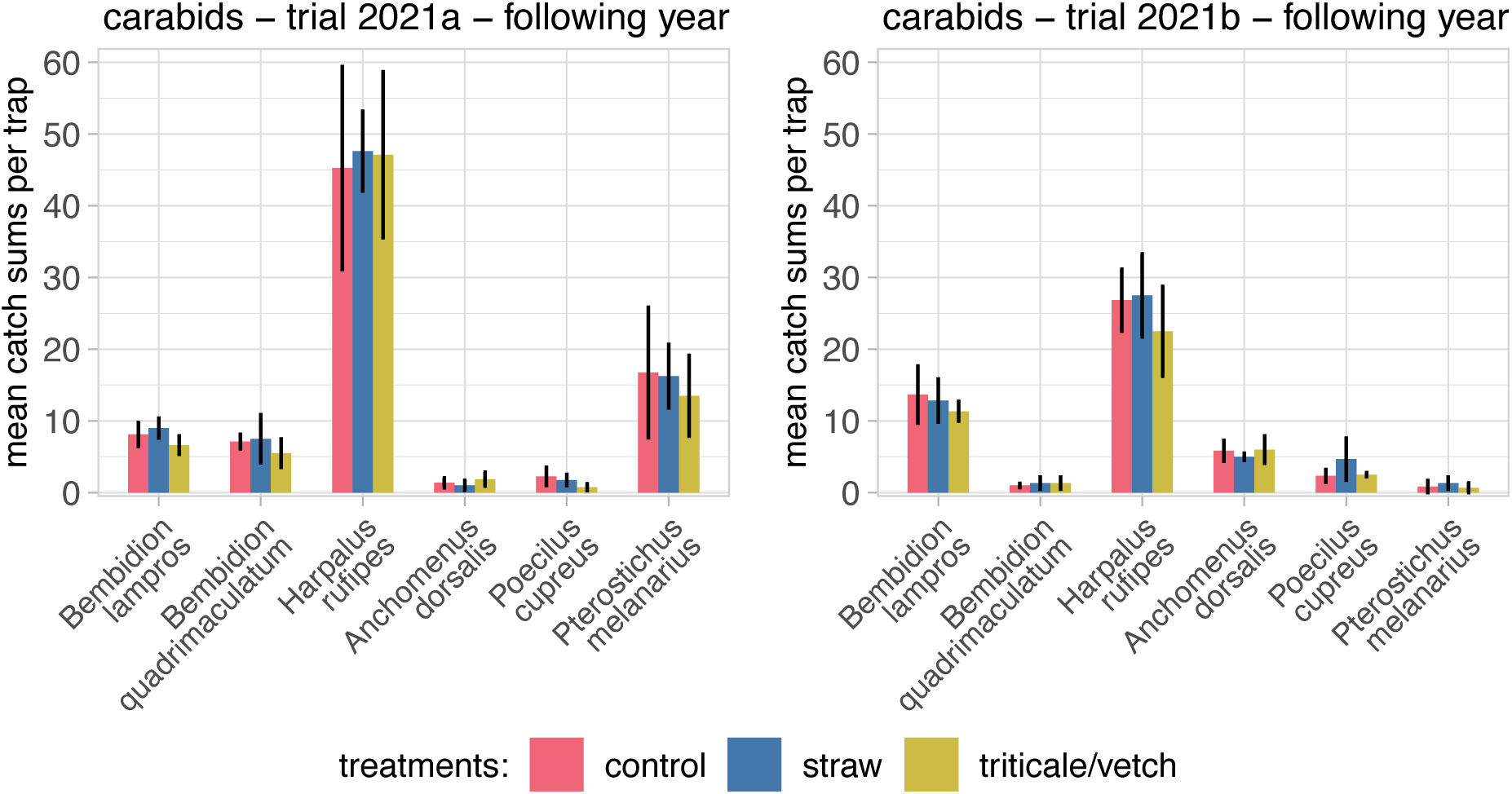
Mean catches (± SE) of the six most frequent carabid species in late July in the year following mulching, with no additional mulching applied in this second year, to investigate potential carry-over eRects. Data are based on ten sampling days per trial. Means and SE were calculated from plot means (n = 4 in trial 2021a and n = 3 in trial 2021b).

### 3.3. Carabid community structure

Shannon diversity and species richness metrics showed no consistent pattern across trials (Figure 6). However, species richness tended to be higher in triticale/vetch plots, particularly in trials 2021a and 2021b, while Shannon diversity was slightly lower in triticale/vetch plots in trial 2021a.

**Figure 6:**
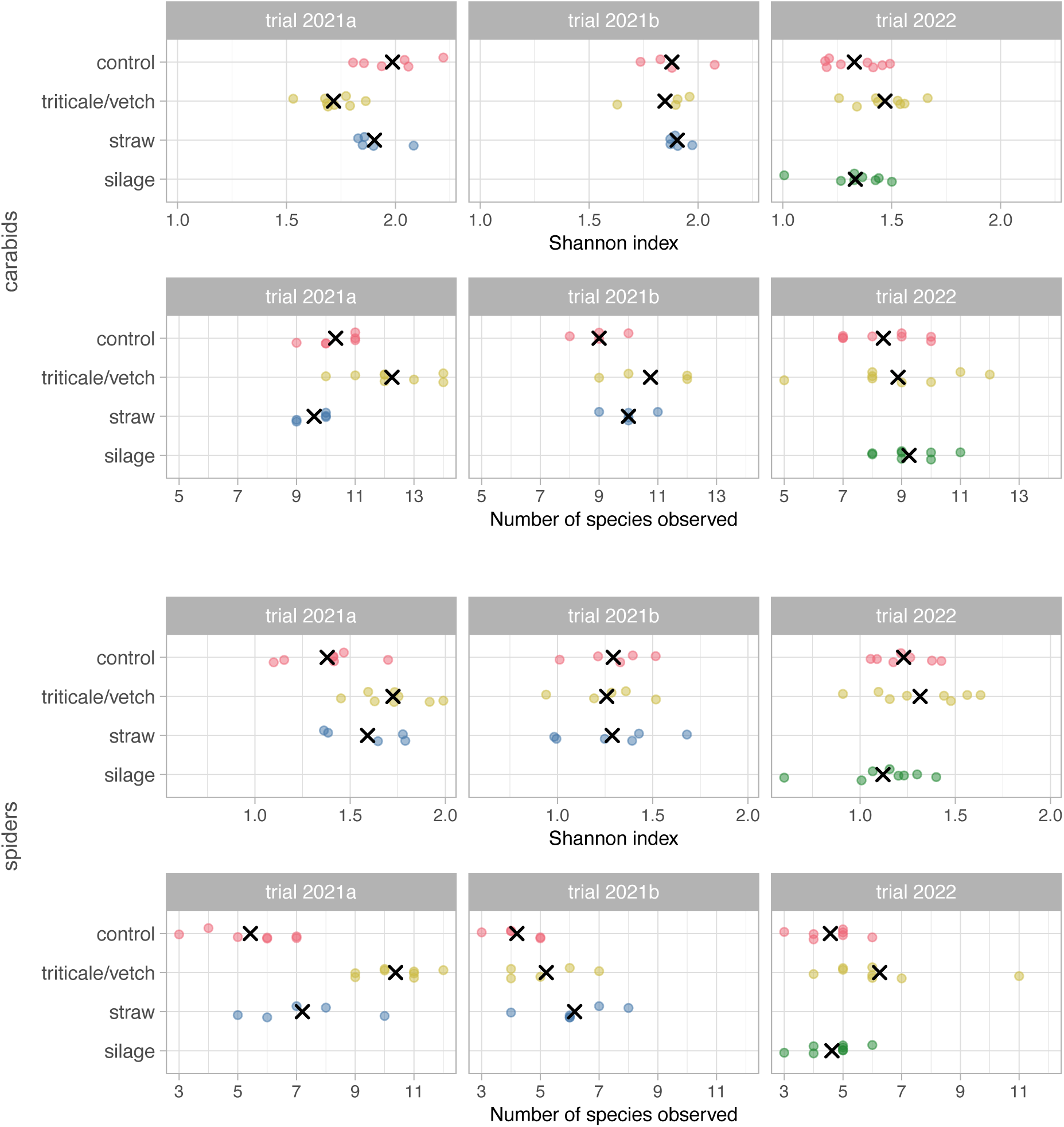
Distribution of species numbers and evenness within the trials and treatments for carabids and spiders. X indicate mean values.

NMDS analysis revealed significant treatment effects on community composition in trials 2021a and 2022 (Figure 7). In trial 2021a, triticale/vetch plots clearly separated from other treatments along NMDS1 (PERMANOVA: p=0.001, pseudo-F=6.81), with significant differences between control-triticale/vetch (p=0.012) and control-straw (p=0.018) pairs. In trial 2022, community structure in control plots differed significantly from both mulch treatments (PERMANOVA: p=0.001, pseudo-F=6.69), with pairwise differences between control-triticale/vetch and control-silage pairs (both p=0.006).

**Figure 7:**
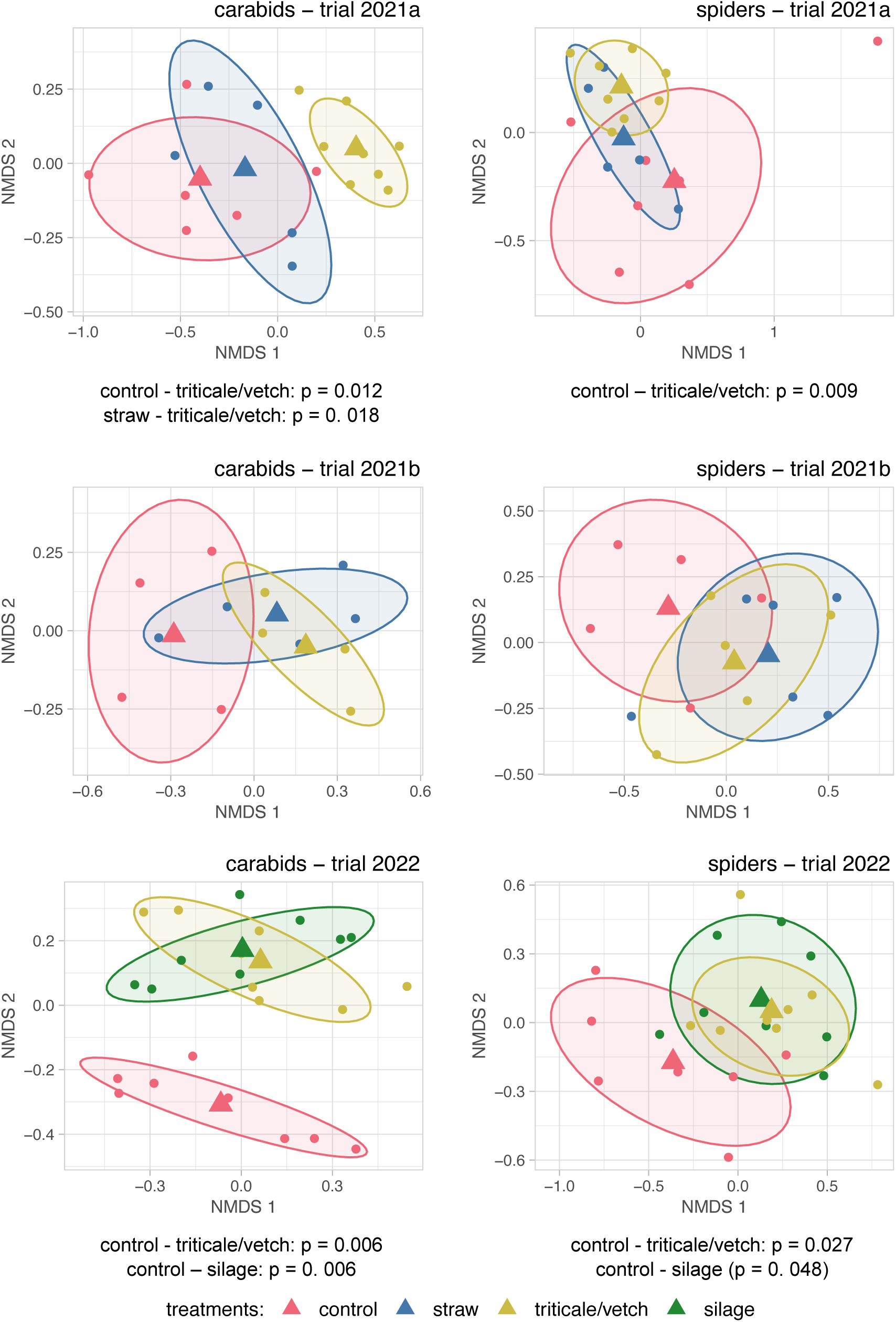
Non-metric multidimensional scaling (NMDS) ordinations performed on the Bray-Curtis dissimilarity matrices calculated from carabid and spider abundance data. Ellipses encompassing 70% of the points with centroids are shown for each treatment group to facilitate visual assessment. Stress values of the ordinations were A: 0.061, B: 0.095, C: 0.141 for carabid data and A: 0.091, B: 0.109, C: 0.102 for spider data. Significant test results from pairwise PERMANOVAs with Bonferroni correction are shown below the ordination plots.

### 3.4. Spider abundances

Spider activity densities were consistently higher in July than in June across all trials (Figure 3). The triticale/vetch treatment led to marked increases in spider abundance, with 2.2-, 1.2-, and 2.2-fold higher captures compared to control plots in trials 2021a, 2021b, and 2022, respectively. Straw and silage treatments also resulted in moderately elevated spider numbers, ranging from 1.3 to 1.6 times the control.

The six most frequently recorded spider taxa were the five Linyphiidae species *Erigone dentipalpis*, *Oedothorax apicatus*, *Erigone atra*, *Tenuiphantes tenuis*, *Agyneta rurestris*, and the lycosid *Pardosa agrestis* (see Appendix 2 for a detailed species list). *E. dentipalpis* and *O. apicatus* dominated across trials (Figure 4), but their responses to treatments diverged:

- *E. dentipalpis* showed consistently positive responses to triticale/vetch (1.4-3.7 times higher than control) and straw treatments (1.9-2.3 times higher).
- *E. atra* showed similar but less pronounced increases in mulched plots (1.5-2.8 times higher in trials 2021a and 2021b).
- *O. apicatus*, however, exhibited no significant response to any mulch treatment.
- *T. tenuis* was more abundant in mulched plots in trial 2021a but showed the opposite trend in other trials.
- *P. agrestis* benefited from triticale/vetch mulching in trials 2021a and 2022 but not in trial 2021b.

The multi-level pattern analysis showed no significant associations for either trial.

#### 3.3.1 Carry-over eDects

Sampling in September and post-harvest sampling in October of trial 2021a almost exclusively captured *O. apicatus*, with 1.7–2.0 times higher numbers in straw and triticale/vetch treatments compared to control in October (Appendix 3).

#### 3.3.2 Spider community structure

Spider communities showed similar Shannon diversity across treatments in all trials, with no clear pattern (Figure 6). However, species richness tended to be higher in both mulched treatments compared to control.

NMDS analysis showed significant separation of spider communities between treatments in trials 2021a and 2022 (Figure 7). In trial 2021a, communities separated primarily along NMDS2 (PERMANOVA: p=0.004, pseudo-F=2.33), with significant differences between triticale/vetch and control (p=0.009). In trial 2022, both mulch treatments separated from control (PERMANOVA: p=0.011, pseudo-F=2.79), with significant pairwise differences (control-triticale/vetch: p=0.027; control-silage: p=0.048).

### 3.4. Mulch-activity bias

No consistent differences were found in catches between pitfall traps surrounded by mulch versus those with a cleared area for either carabids or spiders (Appendix 4). However, slightly higher catches of A. dorsalis were observed in pitfall traps with cleared surroundings in triticale/vetch plots, and O. apicatus showed higher captures in cleared areas within triticale/vetch plots in trial 2021a. Conversely, both Erigone species were more frequently captured in traps surrounded by straw mulch within straw plots of trial 2021b.

## 4 Discussion

Our study demonstrates that organic mulching significantly influences ground-dwelling predator communities in potato agroecosystems, with effects varying by mulch type and arthropod species (i). Triticale/vetch mulch consistently showed the strongest impact, substantially increasing the abundance of specific carabid species – particularly *P. cupreus* (3.3–9.0 times control), *B. lampros*, and *B. quadrimaculatum* – and several Linyphiidae spider species, especially *E. dentipalpis* (1.4–3.7 times control) and *E. atra*. These abundance changes were accompanied by significant shifts in overall community composition, as confirmed by NMDS ordinations and PERMANOVA analyses, while species richness tended to be higher in mulched plots.

Sampling beyond the main growing season (i.e., for carry-over effects) indicated that the positive impacts of mulching on predator communities were generally limited in their seasonal persistence (ii). For instance, elevated abundances of *H. rufipes* were detected in mulched plots in September in trial 2021a, and *O. apicatus* remained moderately more abundant in mulched plots until October. However, these effects did not carry over into the following year for carabid communities. The absence of such long-term carry-over implies that annual mulch application is likely necessary to sustain benefits for these predators. This lack of persistence into the subsequent year can be attributed to several interconnected mechanisms: first, intensive soil disturbance during potato harvest disrupts the mulch-induced habitat structures; second, carabid beetles exhibit pronounced seasonal migration patterns, leaving the field centres in autumn to overwinter in field margins and adjacent semi-natural habitats (Knapp et al., 2019; Wallin, 1985, 1988); and third, in spring, these predators recolonize the field from overwintering sites, with recolonization supposedly occurring independently of the previous year’s mulch treatment. Consequently, mulching in potato cultivation should be viewed as a recurrent, active management measure rather than a one-oE, long-term habitat improvement.

### 4.1. Implications for pest regulation

The enhancement of carabid beetle and spider populations by organic mulching has direct implications for biological pest control. Both groups are well-documented predators of aphids (Chiverton, 1987; Nyffeler & Benz, 1988b, 1988a; K. D. Sunderland & Vickerman, 1980), and some carabid species are known to prey on the Colorado potato beetle (*Leptinotarsa decemlineata*) (Chang & Snyder, 2004; Hazzard et al., 1991). Two studies conducted in parallel to the present work – one fully (Winkler et al., 2024) and one partly (Winkler et al., 2025) utilizing the same field trials – demonstrated that mulching led to marked reductions in Colorado potato beetle and aphid populations. While multiple mechanisms were discussed as contributing to these pest reductions, the increased presence of carabid and spider predators likely played an additional role. Thus, our findings support the concept that habitat management practices such as organic mulching can contribute to conservation biological control by promoting predator populations and enhancing pest suppression.

### 4.2. Drivers of mulch eDects

Three primary mechanisms likely explain the observed effects of organic mulching on predator communities. First, mulch increases **habitat complexity**, providing diverse microstructures that benefit particular species based on their ecological traits. Web-building Linyphiidae spiders (*E. dentipalpis, E. atra, A. rurestris*) likely benefited from additional anchoring points for web construction in the otherwise structurally simple potato field environment. This interpretation aligns with previous research showing that wheat stems, and cereal stubble improve web construction by *T. tenuis* (Harper, 2020; K. D. Sunderland et al., 1986) and that complex habitats generally increase Linyphiidae abundance (Bell et al., 2002; Birkhofer et al., 2008). On the other hand, organic mulching may impede the movement of arthropods, potentially affecting both prey and predator species (Szendrei et al., 2009; Weiler et al., 2025b). In other ecosystems, increased structural complexity has been shown to reduce intraguild predation by providing spatial refuges that limit antagonistic interactions among predator species (Finke & Denno, 2002).

Second, mulching creates favourable **microclimate conditions** by increasing moisture retention, reducing temperature extremes, and providing shade (Brown & Gallandt, 2018; Dudás et al., 2016; Edwards et al., 2000; Johnson et al., 2004). These conditions may benefit diurnal, spring-breeding carabids, such as *P. cupreus*, *B. lampros*, and *B. quadrimaculatum*, more than nocturnal, autumn-breeding species, such as *Pt. melanarius* and *H. rufipes*, as they may be more sensitive to desiccation and temperature fluctuations. However, during the extreme drought and high temperatures observed in 2022 (see Figure 2), the buffering effect of mulch appeared insufficient to fully mitigate macroclimatic stress, resulting in less pronounced or altered responses among arthropods. For example, while *P. cupreus* consistently benefited from mulching, whereas *Bembidion* spp. showed reduced abundance in the dry year and *A. dorsalis* even preferred control plots. This highlights that the habitat value of mulch is context-dependent and varies with species-specific stress tolerances and ecological niches.

Third, mulch further alters **prey availability** in complex ways. While reducing pest densities such as aphids (Saucke & Döring, 2004; Winkler et al., 2025) and Colorado potato beetles (Junge et al., 2022; Winkler et al., 2024), mulch simultaneously enhances soil microbial and fungal activity, indicating stimulation of the detrital food web (Henzel et al., 2025), and promotes alternative prey like *Collembola* (Birkhofer et al., 2008; Sereda et al., 2015). By increasing the abundance of decomposers and detritivores, mulching can provide a more stable and diverse prey base for generalist predators, especially during periods of low pest density. This resource subsidy from the below- ground food web supports higher predator densities and persistence in the field, thereby strengthening top-down control of herbivores when pest populations rise (Scheu, 2001). Predator responses were likely influenced by species-specific dietary preferences, with some species benefiting more from increased detritivore prey, while others were more affected by reduced pest availability (Bilde et al., 2000; Bilde & Toft, 2001; Kielty et al., 1999; Mundy et al., 2000).

### 4.3. Comparison with previous research

Our findings both support and extend previous studies on organic mulching effects on beneficial arthropods. The observed increase in carabid abundance aligns with North American research by Johnson et al. (2004) and Brust (1994), who found higher carabid numbers in straw-mulched potatoes. However, our results contrast with Schmidt et al. (2004), who reported no differences in carabid abundance in straw-mulched cereals, and Rämert (1996), who found higher *Bembidion* populations in unmulched carrot plots.

These discrepancies likely reflect differences in crop systems, regional arthropod communities, and experimental methodologies.

For spiders, our findings corroborate previous studies showing that Linyphiidae benefit from organic mulching. Johnson et al. (2004) and Schmidt et al. (2004) reported more Linyphiidae in straw-mulched potatoes and cereals, respectively. Similarly, Halaj et al. (2000) observed dramatic increases in spider numbers in straw refugia within soybean fields. The positive response of *Erigone* species to reduced tillage with cereal stubble reported by Volkmar et al. (2003) further supports our findings.

Our study advances the research field by providing species-level identification of predator responses across multiple mulch types and connecting responses to species-specific ecological traits.

## 5 Strengths and limitations

Our study possesses several notable strengths. By spanning two years and three field sites, we increased the validity of our findings, extending their relevance beyond a single location or season. The experimental design included adequate replication and large plot sizes, and the sampling periods were sufficient to capture key aspects of the seasonal dynamics of ground-dwelling arthropod communities. Furthermore, the identification of carabids and spiders to species level facilitated detailed community analyses and provided species-specific insights.

Potential sampling biases were addressed by comparing pitfall trap catches from traps surrounded by mulch to those from traps with mulch cleared within a 1-m radius. This comparison revealed no consistent differences in total carabid or spider catches; however, certain species (e.g., *A. dorsalis*, *O. apicatus*, *Erigone* spp.) exhibited minor, species-specific variations. These findings suggest that our main conclusions are robust, although some imprecision for individual species cannot be entirely excluded.

Nonetheless, our study was limited to central Germany and focused exclusively on potato cultivation, which may constrain the generalizability of our results to other regions or cropping systems. As with all pitfall trapping studies, we measured activity-density rather than true abundance (Luff, 1975; Topping & Sunderland, 1992). Still, the comparability of methods and the stable patterns observed across treatments provide a solid foundation for our conclusions.

## 6 Future perspectives

Building on the results generated in this study, future research should focus on deepening the understanding of the mechanisms by which organic mulching shapes predator communities and ecosystem services in different agroecological contexts. In particular, comparative studies across diverse crop systems and climatic regions would clarify the generalizability of observed patterns and help refine best-practice recommendations for integrated pest management. Further, integrating complementary sampling methods – such as molecular gut content analysis, direct behavioural observations or mark and recapture – could provide more detailed insights into predator-prey interactions and the functional roles of key species promoted by mulching. Long-term field trials, especially those spanning entire crop rotations, will be valuable for assessing the persistence and cumulative effects of repeated mulching on both biodiversity and crop productivity.

## 7 Conclusion

The application of organic mulch in potato cultivation offers a straightforward and effective approach to promoting beneficial arthropod diversity at a time when insect declines are a major concern. By increasing habitat complexity and moderating the microclimate, mulching supports key predator species in pest regulation. This ecological service is especially valuable as agriculture seeks alternatives to insecticides.

Beyond arthropod conservation, mulching offers additional agronomic benefits. Improved soil structure, increased water retention, enhanced microbial activity, and effective weed suppression collectively promote crop health and support more sustainable yields. Our results indicate that integrating organic mulch into crop management can deliver both ecological and economic advantages, offering farmers a feasible means to balance productivity with conservation objectives.

## Acknowledgments

We thank Rainer Wedemeyer and Joachim Deckers, as well as the farmers for managing the field trials. We would also like to thank Gero Jäger for his advice on spider identification and are grateful to Inger Deilke, Pauline Reichardt and Lara Schubert for assisting with arthropod sampling and identification.

## Appendix

### 1.1 Appendix 1

Dates of trap emptying of and sampling days for trials 2021a, 2021b and 2022. Carabids were identified on all dates, spiders on the dates printed in bold. Dashed lines between dates indicate a period of no sampling in between these dates.

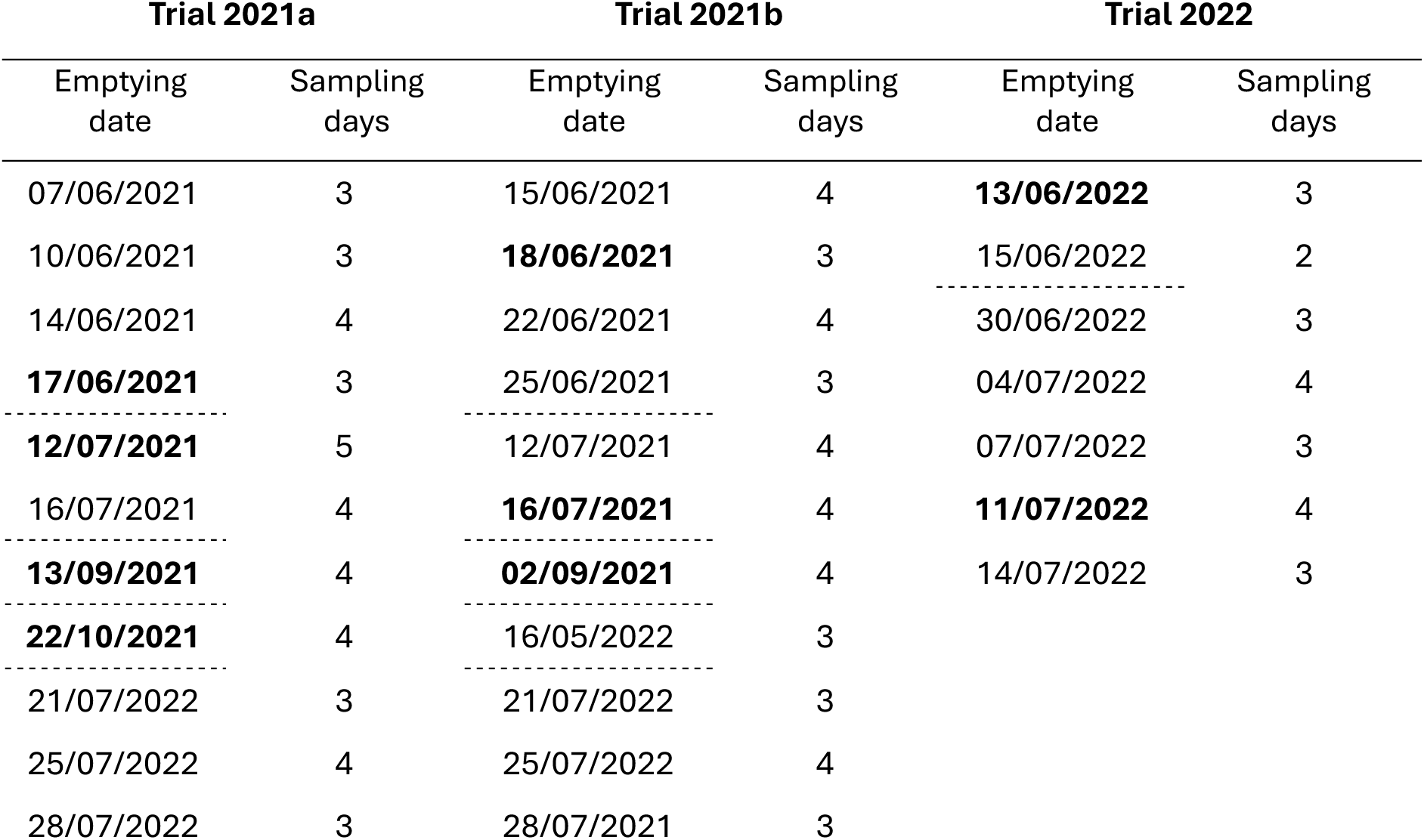

| Trial 2021a |  | Trial 2021b |  | Trial 2022 |  |
| --- | --- | --- | --- | --- | --- |
| Emptying date | Sampling days | Emptying date | Sampling days | Emptying date | Sampling days |
| 07/06/2021 | 3 | 15/06/2021 | 4 | **13/06/2022** | 3 |
| 10/06/2021 | 3 | **18/06/2021** | 3 | 15/06/2022 | 2 |
| 14/06/2021 | 4 | 22/06/2021 | 4 | 30/06/2022 | 3 |
| **17/06/2021** | 3 | 25/06/2021 | 3 | 04/07/2022 | 4 |
| **12/07/2021** | 5 | 12/07/2021 | 4 | 07/07/2022 | 3 |
| 16/07/2021 | 4 | **16/07/2021** | 4 | **11/07/2022** | 4 |
| **13/09/2021** | 4 | **02/09/2021** | 4 | 14/07/2022 | 3 |
| **22/10/2021** | 4 | 16/05/2022 | 3 |  |  |
| 21/07/2022 | 3 | 21/07/2022 | 3 |  |  |
| 25/07/2022 | 4 | 25/07/2022 | 4 |  |  |
| 28/07/2022 | 3 | 28/07/2021 | 3 |  |  |

### 1.2 Appendix 2

Species of carabid beetles and spiders identified in the three trials in order of abundance.

9.2 Appendix 2 *Species of carabid beetles and spiders identified in the three trials in order of abundance.*
| **Carabids** |  | **Spiders** |  |
| --- | --- | --- | --- |
| Species | abundance | Species | abundance |
| *Harpalus rufipes* | 2,613 | *Erigone dentipalpis* | 754 |
| *Bembidion lampros* | 2,507 | *Oedothorax apicatus* | 721 |
| *Poecilus cupreus* | 1,906 | *Erigone atra* | 164 |
| *Bembidion quadrimaculatum* | 1,042 | *Tenuiphantes tenuis* | 126 |
| *Anchomenus dorsalis* | 1,003 | *Agyneta rurestris* | 119 |
| *Pterostichus melanarius* | 772 | *Pardosa agrestis* | 46 |
| *Harpalus affinis* | 596 | *Bathyphantes gracilis* | 39 |
| *Nebria brevicollis* | 447 | *Trochosa ruricola* | 19 |
| *Trechus quatristriatus* | 427 | *Pardosa spp.* | 14 |
| *Microlestes minutulus* | 228 | *Pachygnatha degeeri* | 11 |
| *Loricera pillicornis* | 177 | *Araeoncus humilis* | 10 |
| *Zabrus tenebrioidis* | 58 | *Oedothorax retusus* | 9 |
| *Amara similata* | 41 | *Pardosa palustris* | 8 |
| *Amara familiaris* | 40 | *Oedothorax agrestis* | 8 |
| *Calathus fuscipes* | 37 | *Drassyllus spp.* | 7 |
| *Amara consularis* | 26 | *Gnaphosidae spp.* | 7 |
| *Bembidion properans* | 20 | *Porrhomma spp.* | 6 |
| *Asaphidion flavipes* | 19 | *Diplostyla concolor* | 5 |
| *Notiophilus biguttatus* | 19 | *Micaria spp.* | 3 |
| *Bembidion tetracolum* | 15 | *Pardosa amentata* | 3 |
| *Amara aenea* | 14 | *Pelecopsis parallela* | 3 |
| *Acupalpus meridianus* | 9 | *Histocona torpida* | 2 |
| *Bembidion obtusum* | 8 | *Pardosa agricola* | 2 |
| *Clivina fossor* | 8 | *Phrurolithus minimus* | 2 |
| *Demetrias atricapillus* | 5 | *Xysticus cochi* | 2 |
| *Pterostichus niger* | 5 | *Xysticus spp.* | 2 |
| *Brachinus crepitans* | 4 | *Histocona torpida* | 2 |
| *Brachinus explodens* | 4 | *Tibellus oblungus* | 1 |
| *Calathus erratus* | 3 | *Callilepis nocturna* | 1 |

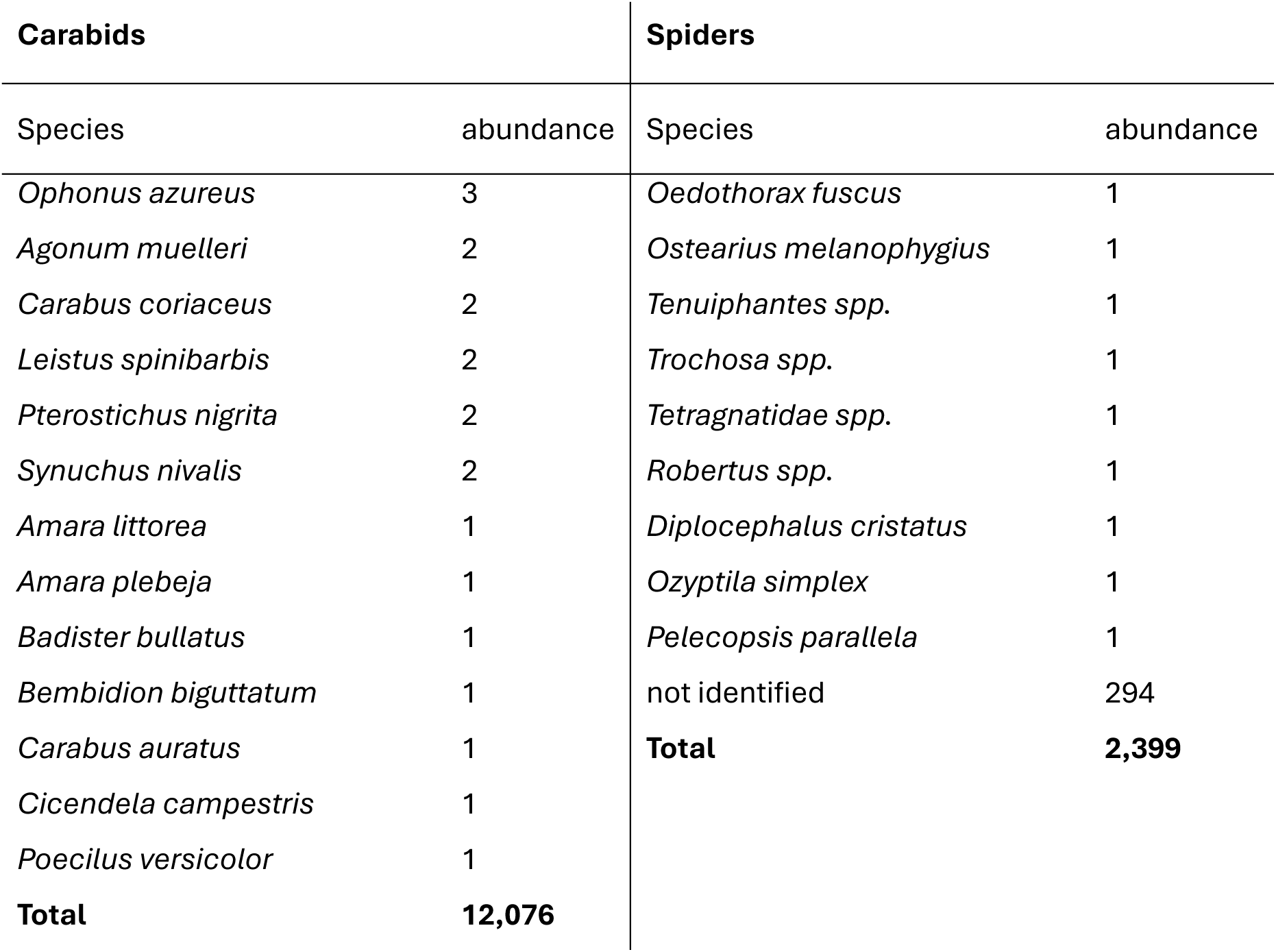

| Carabids |  | Spiders |  |
| --- | --- | --- | --- |
| Species | abundance | Species | abundance |
| *Ophonus azureus* | 3 | *Oedothorax fuscus* | 1 |
| *Agonum muelleri* | 2 | *Ostearius melanophygius* | 1 |
| *Carabus coriaceus* | 2 | *Tenuiphantes* spp. | 1 |
| *Leistus spinibarbis* | 2 | *Trochosa* spp. | 1 |
| *Pterostichus nigrita* | 2 | *Tetragnatidae* spp. | 1 |
| *Synuchus nivalis* | 2 | *Robertus* spp. | 1 |
| *Amara littorea* | 1 | *Diplocephalus cristatus* | 1 |
| *Amara plebeja* | 1 | *Ozyptila simplex* | 1 |
| *Badister bullatus* | 1 | *Pelecopsis parallela* | 1 |
| *Bembidion biguttatum* | 1 | not identified | 294 |
| *Carabus auratus* | 1 | **Total** | **2,399** |
| *Cicendela campestris* | 1 |  |  |
| *Poecilus versicolor* | 1 |  |  |
| **Total** | **12,076** |  |  |

### 1.3 Appendix 3

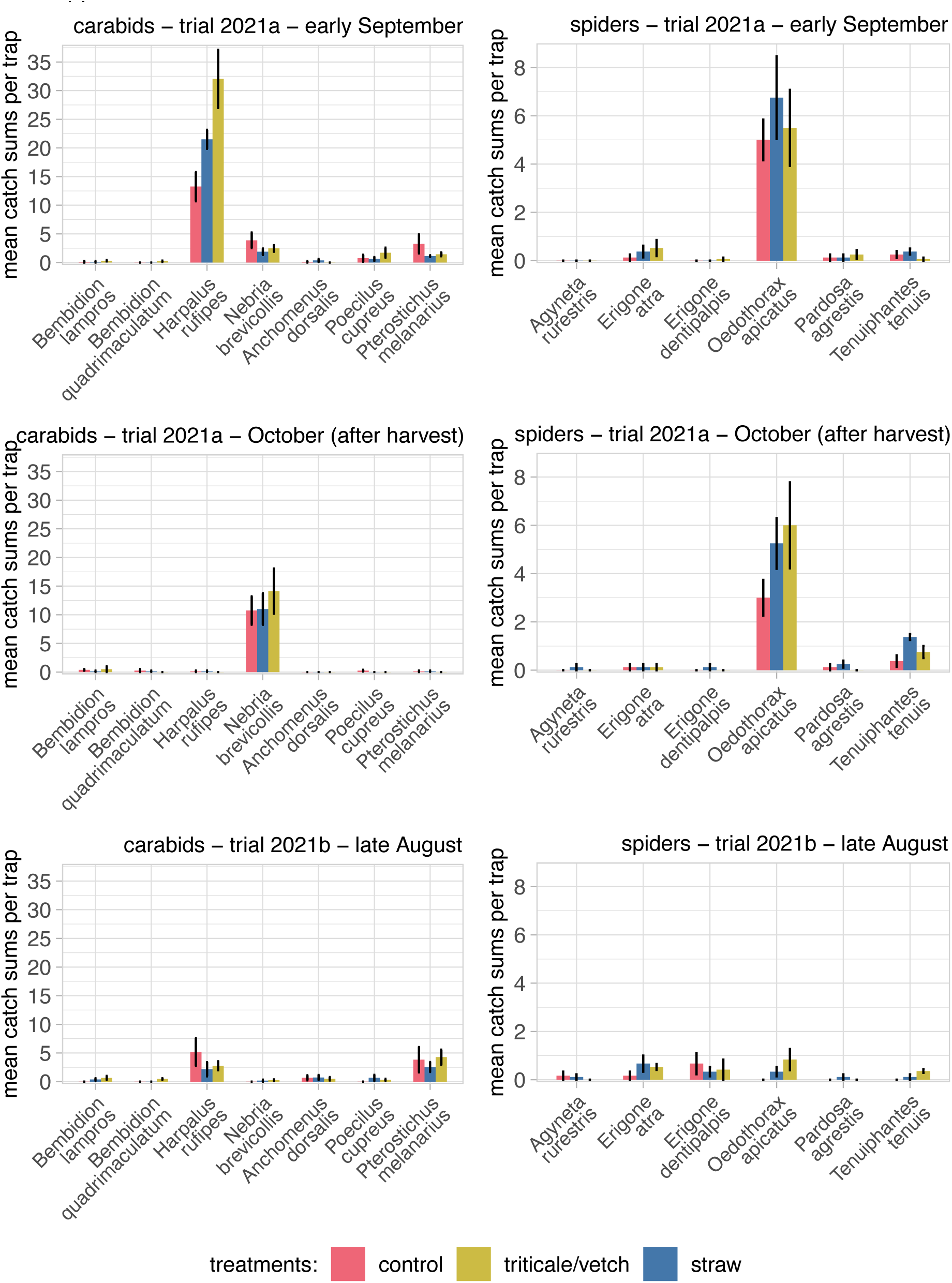

Mean catches (± SE) of the seven most frequent carabid species in late August to late October from four sampling days each. Values represent means and SE calculated from plot means (n = 4 in trial 2021a and n = 3 in trial 2021b).

### 1.4 Appendix 4

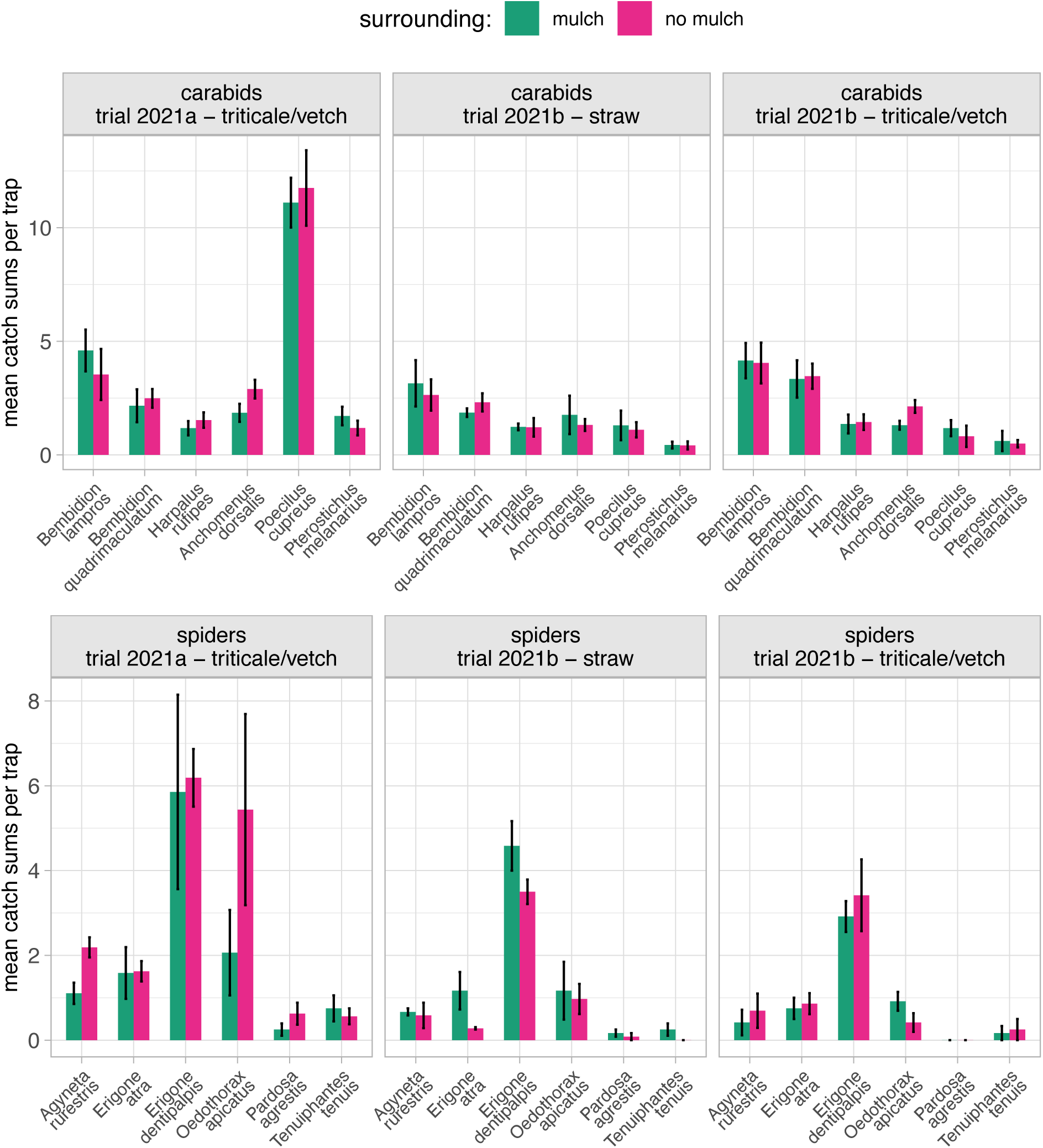

Comparison of mean catches (± SE) of traps surrounded by mulch or no mulch. The summarised catches from June/July of the six most common carabid and spider species are shown. Values represent means and SE calculated from plot means (n = 4 in trial 2021a and n = 3 in trial 2021b).

